# Perceptual Degradation Affects Stop-Signal Performance in Normal Healthy Adults

**DOI:** 10.1101/2020.10.29.351502

**Authors:** Maria V. Soloveva, Sharna D. Jamadar, Matthew Hughes, Dennis Velakoulis, Govinda Poudel, Nellie Georgiou-Karistianis

## Abstract

During stop-signal task performance, little is known how the quality of visual information of the ‘go’ stimuli may indirectly affect the interplay between the ‘go’ and ‘stop’ processes. In this study, we assessed how perceptual degradation of the visual ‘go’ stimuli affect response inhibition. Twenty-six healthy individuals (mean age 33.34 ± 9.61) completed a modified 12-minute stop-signal task, where ‘V’ and ‘Y’ letters were used as visual ‘go’ stimuli. The stimuli were subjected to four levels of perceptual degradation using Gaussian smoothing, to parametrically manipulate stop difficulty across low, intermediate-1, intermediate-2 and high difficulty conditions. On 33% of trials, the stop-signal (50ms audio tone) followed a ‘go’ stimulus after a stop-signal delay, which was individually adjusted for each participant. As predicted, we found that with increased level of stop difficulty (little perceptual degradation), reaction times on ‘go’ trials and the proportion of successful behavioural inhibitions on ‘stop’ trials (*P*(i)) decreased in normal healthy adults. Contrary to our predictions, there was no effect of increased stop difficulty on the number of correct responses on ‘go’ trials and reaction times on ‘stop’ trials. Overall, manipulation of the completion time of the ‘go’ process via perceptual degradation has been partially successful, whereby increased stop difficulty differentially affected *P*(i) and SSRT. These findings have implications for the relationship between the ‘go’ and ‘stop’ processes and the horse-race model, which may be limited in explaining the role of various cortico-basal ganglia loops in modulation of response inhibition.

**Highlights:**

- Manipulation of the completion time of the ‘go’ process is partially successful
- Perceptual degradation differentially affects stop-signal performance
- Increased stop difficulty (easy ‘go’) results in lower *P*(i)
- Increased stop difficulty (easy ‘go’) has no effect on SSRT
- Horse-race model does not fully explain basal ganglia involvement in inhibition

## Introduction

Response inhibition is a core component of executive function (Barkley 1997; Verbruggen & Logan, 2008), which refers to the ability of an individual to inhibit an unwanted decision, thought, or an ongoing movement in response to unexpected events (Verbruggen & Logan, 2009).

The stop-signal task (SST; Logan & Cowan, 1984) is used to investigate response inhibition in non-clinical and clinical populations. Many studies have documented deficient stop-signal performance in health (Coxon et al., 2012) and disease (Lipszyc & Schachar, 2010), including Parkinson’s disease (Baglio et al., 2011), obsessive-compulsive disorder (OCD; de Wit et al., 2012), schizophrenia (Hughes, Fulham, Johnston, & Michie, 2012), attention deficit hyperactivity disorder (ADHD; Christ, Holt, White, & Green, 2007), Huntington’s disease (Dominguez et al., 2017; Soloveva et al., 2020), as well as normal healthy ageing (Coxon et al., 2012).

In the standard stop-signal paradigm, participants are required to respond rapidly to frequent presentations of ‘go’ stimuli (the ‘go’ task) and on occasional trials inhibit the ‘go’ task response on ‘stop’ trials when a stop-signal is presented just after a ‘go’ stimulus (the ‘stop’ signal task; see Figure 1). Stop-signal task performance has been modelled as the outcome of a race between ‘go’ and ‘stop’ processes, where the winner determines whether response inhibition is successful or unsuccessful (Logan, 1994; Verbruggen & Logan, 2009).

**Fig. 1.**
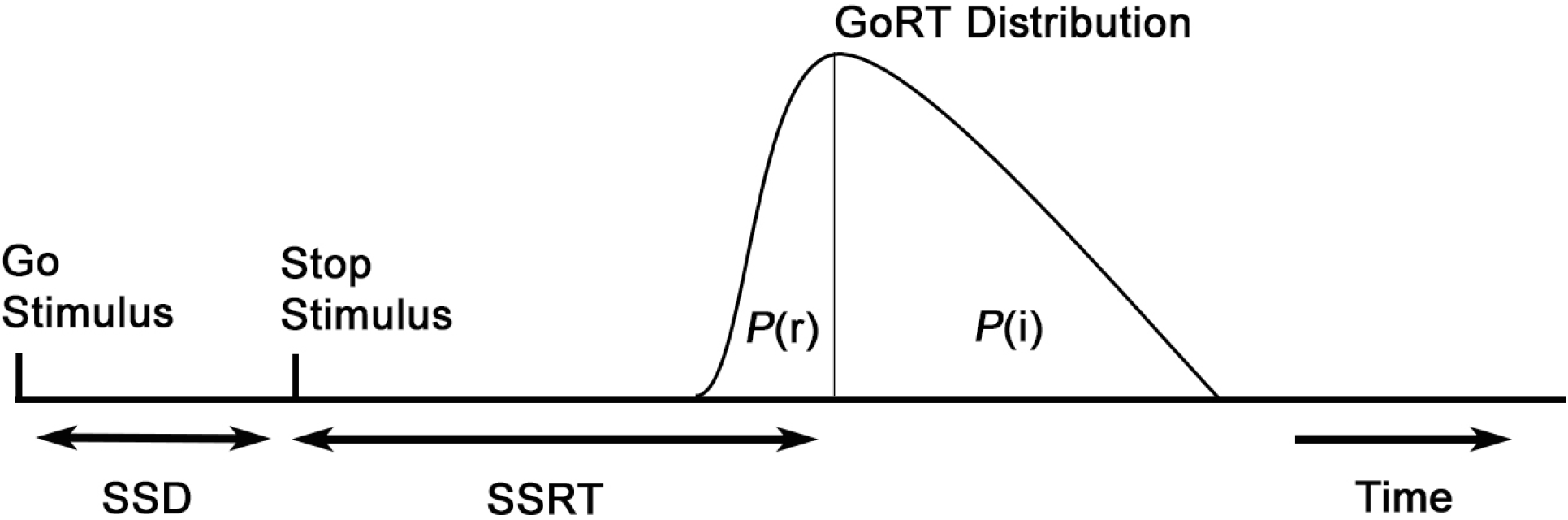
The horse-race model (adapted from Logan & Cowan, 1984). *P*(i) (%) is the probability of successfully inhibiting an initiated motor response and stop-signal reaction time (SSRT) (ms) is the time required to inhibit a motor response – two primary outcome measures of response inhibition in a stop-signal task. Stop-signal delay (SSD) (ms) is the delay between the onset of ‘go’ and stop-signal stimuli. The model suggests that the probability of responding (*P*(r)) and the probability of inhibiting a response (*P*(i)) depends on the distribution of ‘go’ reaction times, SSD and SSRT. Response inhibition is determined by the outcome of a race between two distinct processes – a ‘go’ process and a ‘stop’ process. A faster ‘go’ process results in a high probability of responding on a stop-signal trial (*P*(r), unsuccessful inhibition). Slow ‘go’ process results in a low probability of responding on a stop-signal trial (*P*(i), successful inhibition). SSRT is computed from the distribution of correct RTs (ms) on ‘go’ trials and the proportion of successful inhibitions on ‘stop’ trials

The most direct method to manipulate task difficulty in the stop-signal task is to vary the stop-signal delay (SSD) duration (D’Alberto et al., 2018; Hughes, Johnston, Fulham, Budd, & Michie, 2013; Verbruggen et al., 2019). Consistent with the horse-race model, if SSD is short, response inhibition is successful (i.e., all ‘go’ reaction times are inhibited, *P*(i) = 1)); however, as SSD increases, *P*(i) decreases, until SSD is long enough for all ‘go’ reaction times to escape inhibition (*P*(i) = 0) (Hughes et al., 2013). At shorter SSDs, where the stop signal appears very soon after the ‘go’ stimulus is presented, individuals are more likely to suppress motor response activation; whereas if SSD is longer, motor response activation is more likely to carry on until completion. Importantly, increasing or decreasing SSD to manipulate stopping difficulty fails to account for inter-individual variability in reaction times on ‘go’ trials (Hughes et al., 2013). Specifically, to increase the number of successful behavioural stops (*P*(i)), individuals may strategically increase reaction times on ‘go’ trials, when they expect a stop-signal to occur and decrease reaction times on ‘go’ trials when not expecting (Leotti & Wager, 2009; Verbruggen & Logan, 2009). D’Alberto et al. (2018) showed that differences between superior (mean SSRT = 137 ms) and poor (SSRT = 250 ms) inhibitors, in stop-related functional brain activity in the response inhibition network, were attenuated when controlling for SSD duration (i.e., equal stop difficulty for both groups). Taken together, the SSD that predicts a given *P*(i) in one individual may elicit a completely different *P*(i) in another, attenuating the effect of task difficulty on inhibitory processes at both the behavioural and brain levels (D’Alberto et al., 2018; Hughes et al., 2013; Leotti & Wager, 2009).

The horse-race model shows similarities to the proposed functions of the cortico-basal ganglia loops for motor control (Freeze, Kravitz, Hammack, Berke, & Kreitzer, 2013; Mallet et al., 2016). The striatum and subthalamic nucleus are the input stations of the basal ganglia that converge on substantia nigra pars reticulata, which determines behavioural outcomes - motor initiation or inhibition (see Figure 2) (Jahanshahi, Obeso, Rothwell, & Obeso, 2015; Schmidt & Berke, 2017). It is known that activation of the direct ‘go’ pathway tends to promote movement while activation of the indirect ‘stop’ pathway suppresses movement, suggesting that response inhibition is modelled as the outcome of a race between ‘go’ and ‘stop’ neural pathways in basal ganglia (Albin et al., 1989; Aron & Poldrack, 2006; Freeze et al., 2013; Schmidt & Berke, 2017; Schmidt, Leventhal, Mallet, Chen, & Berke, 2013; Wei & Wang, 2016). Furthermore, death of neurons within these striatal pathways affects movement planning and execution, particularly in neurodegenerative disorders, such as Huntington’s and Parkinson’s disease (Jamadar, 2019; Poudel et al., 2019).

**Fig. 2.**
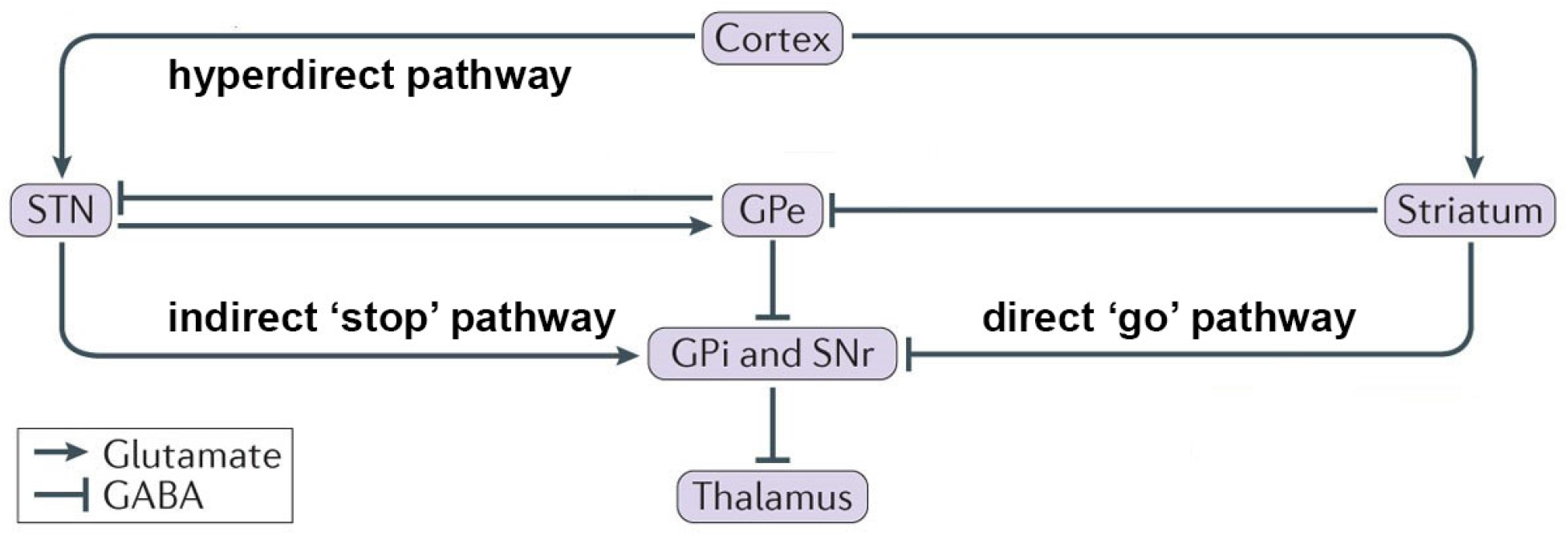
Visual representation of the cortico-basal ganglia circuitry, which is implicated in motor control (adapted from Jahanshahi et al., 2015). This circuitry is comprised of three neural pathways, including a direct pathway that promotes movement, an indirect pathway that suppresses motor behavior, and a hyperdirect pathway thought to suppress inappropriate movements. Striatal neuronal projections to substantia nigra pars reticulata (SNr) correspond to a direct ‘go’ neural pathway. Projections from the subthalamic nucleus (STN) to substantia nigra form an indirect ‘stop’ neural pathway. Neuronal output from the striatum and subthalamic nucleus converge on substantia nigra pars reticulata, providing responses to ‘go’ and ‘stop’ processes. When striatal ‘go’ neurons reach substantia nigra first (fast ‘go’ process; striatal inhibition is strong) the ‘go’ process wins, and this results in unsuccessful inhibition; if the ‘go’ process is slow (striatal inhibition is low) and neurons from subthalamic nucleus-based neural pathway reach substantia nigra first – the ‘stop’ process wins, and this results in successful inhibition. The neuronal output then goes through the thalamus and returns to the cortex. The subthalamic nucleus has excitatory and glutamatergic projections to the external segment of the globus pallidus (GPe), the GPi and the dopaminergic nigro-striatal pathway. All other internal connections within the basal ganglia network are GABAergic and inhibitory

Considering the stop-signal task, if the ‘stop’ cue occurs before movement onset, striatal inhibition is low, allowing neuronal output from subthalamic nucleus to reach substantial nigra, and this results in successful motor inhibition (Hughes, Fulham, & Michie, 2016; Schmidt et al., 2013). By contrast, if the ‘stop’ cue occurs only briefly before movement onset, strong inhibition from striatum prevents the substantia nigra to respond to that cue to inhibit movement (Schmidt et al., 2013). When communication from the striatal ‘go’ neurons reaches substantia nigra first, the ‘go’ process is fast, whereas if communication from the neurons in the subthalamic nucleus reaches substantia nigra first, the ‘go’ process is slow. As such, a fast ‘go’ process can produce unsuccessful inhibition (‘go’ wins) and a slow ‘go’ process results in successful inhibition (‘stop’ wins) (Schmidt et al., 2013; Wei & Wang, 2016). Overall, this suggests that if we manipulate the completion time of the ‘go’ process, then we can indirectly manipulate the difficulty of the ‘stop’ process.

An alternate way to manipulate stop difficulty in the stop-signal paradigm is to make visual ‘go’ stimuli perceptually more challenging, thereby indirectly affecting the ability of individuals to inhibit a motor response. If it takes more time for an individual to process perceptually challenging ‘go’ stimuli, then it should be easier to stop the ‘go’ response after the stop-signal, compared to trials where individuals process ‘go’ stimuli quickly. For example, Jahfari, Ridderinkhof, and Scholte (2013) used images of male and female faces as visual ‘go’ stimuli during stop-signal performance, and the visual stimuli contained either all (hard ‘stop’ trials), low (intermediate), and high spatial frequencies (easy ‘stop’ trials). It was found that at high spatial frequency condition (perceptually challenging ‘go’ stimulus; easy ‘stop’ trials), healthy adults demonstrated increased ‘go’ RTs (ms), confirming that the stimuli were difficult to discriminate; and decreased SSRTs (ms), indicating improved stop-signal performance. This suggests that with *decreased* level of stop difficulty (hard ‘go’), healthy individuals were slower in responding to perceptually challenging visual ‘go’ stimuli and required less time to inhibit a response after the stop-signal. Of note, there was no effect of spatial frequency information on either *P*(r) or *P*(i), indicating that participants did not show a significant change in the number of unsuccessful and successful behavioural responses when ‘the ‘go’ trial was perceptually challenging. Alternatively, in a study by Ma and Yu (2016), the ‘go’ task was a random-dot coherent motion task, and a visual ‘go’ stimulus was subjected to 8, 15, or 85% coherence, meaning the ‘go’ stimulus was noisier (harder to discriminate) if there was little coherence. In contrast to Jahfari et al. (2013), the mean RTs on ‘go’ trials *and P*(r) increased with increased coherence in healthy adults, which indicates that participants were significantly less accurate in suppressing motor responses when a visual ‘go’ stimulus was easy to discriminate (easy ‘go’; hard ‘stop’). Moreover, participants demonstrated improved stop-signal performance (decreased SSRTs (ms)) with *decreased* coherence (hard ‘go’; easy ‘stop’), suggesting rather a complex picture of the relationship between the ‘go’ and ‘stop’ processes.

Perceptual information is highly relevant for adaptive and flexible every-day behaviours; however, a few studies have attempted to examine how perceptual demands affect stop-signal performance with mixed results (i.e., Jahfari et al., 2013; Ma & Yu, 2016). It is therefore important to better understand whether and how manipulation of the completion time of the ‘go’ process modulates *P*(i) and SSRT in the stop-signal paradigm.

The goal of this study was to determine whether parametric manipulation of perceptual load of visually neutral ‘go’ stimuli (letters ‘Y’ and ‘V’) in the stop-signal task is a robust method to parametrically manipulate stop difficulty at the behavioural level in healthy individuals. Our modified stop-signal task comprised of high perceptual load (low stop difficulty), intermediate-2 perceptual load (intermediate-1 stop difficulty), intermediate-1 perceptual load (intermediate-2 stop difficulty), and low perceptual load (high stop difficulty). In order to ensure that the probability of successful inhibition (*P*(i)) was not affected by variability in the SSD length (see D’Alberto et al., 2018; Hughes et al., 2013), but rather manipulated by the perceptual load of visual ‘go’ stimuli, we individually adjusted SSD based on a participant’s response to intermediate-1 level of stop difficulty.

Primary outcome measures included mean RTs (ms) and the mean number of correct responses (%) on ‘go’ and ‘stop’ trials per stop difficulty condition in healthy adults. Of central interest was the mean RTs on ‘go’ trials and the mean number of successful inhibitions (*P*(i)) (%) on ‘stop’ trials which confirms whether parametric manipulation of perceptual load of visual ‘go’ stimuli in the stop-signal task was successful.

We hypothesised that with increased stop difficulty, RTs on ‘go’ trials and the probability of successful behavioural inhibitions on ‘stop’ trials, would decrease. Further, we hypothesised that with increased stop difficulty, the number of correct responses (%) on ‘go’ trials and RTs on ‘stop’ trials, would increase.

## Methods

### Participants

Twenty-six healthy volunteers (mean age 33.34 ± 9.61; range 22-56; male to female ratio 9:17, 34.6%:65.4%) participated in the study after giving informed consent. The study was approved by the Monash University Research Ethics Committee.

### Modified Stop-signal Task

All participants performed the ~12-minute modified stop-signal task (see Figure 3 (B)). Each block started with a 500ms fixation cross, followed by a series of ‘go’ and ‘stop’ trials. ‘Y’ and ‘V’ letters were used as visual stimuli on ‘go’ and ‘stop’ trials. Here, ‘Y’ and ‘V’ letters were used as visual stimuli on ‘go’ and ‘stop’ trials as target stimuli because the printed letters have similar spatial properties that are difficult to differentiate between at higher levels of perceptual degradation. Participants were required to respond as quickly as possible to the letters ‘Y’ and ‘V’ (each letter presented equiprobably) with the right and left index finger. ‘Go’ stimuli were presented for 150ms each and the stimulus onset asynchrony varied from 1500-2250ms with a mean of 1875ms. On 33% of trials, the stop-signal (50ms, 1000Hz auditory tone) followed the ‘go’ trial after a stop signal delay (SSD) (ms) and participants were required to inhibit a response when the tone was presented.

**Fig. 3.**
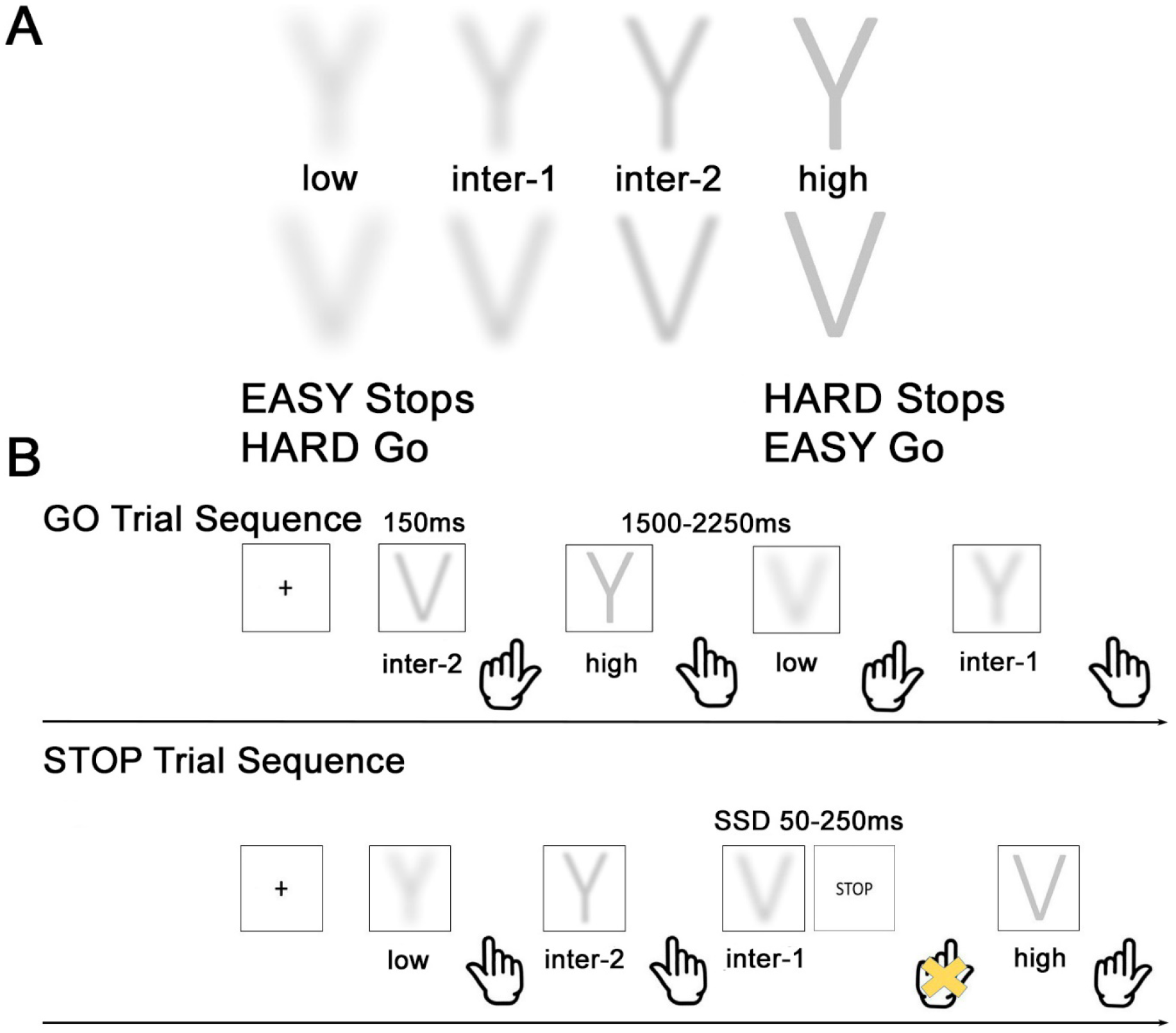
(A) ‘Go’ stimuli (the letters ‘Y’ and ‘V’) subjected to high, intermediate-2, intermediate-1, and low perceptual degradation, corresponding to low, intermediate-1, intermediate-2, and high level of stop difficulty. (B) Visual representation of the modified stop-signal task. In this example, during GO trial sequence, participants responded with the left index finger to identify the letter ‘V’ and with the right index finger to identify the letter ‘Y’ as fast and accurate as they could. During STOP trial sequence, following a ‘go’ trial (‘V’ letter, intermediate-1 level of stop difficulty) a stop-signal was presented, instructing the participant to inhibit the cued response

Importantly, ‘Y’ and ‘V’ letters were subjected to four levels of perceptual degradation using Gaussian smoothing function in MATLAB 8.0 (MATLAB 8.0; http://au.mathworks.com/) to parametrically manipulate stop difficulty (see Figure 3 (A)). The 2-D Gaussian smoothing kernel filter with the standard deviation of 21 corresponded to low level of stop difficulty (easy ‘stop’ trials); a 2-D Gaussian smoothing kernel filter with the standard deviation of 14 corresponded to intermediate-1 level of stop difficulty; a 2-D Gaussian smoothing kernel filter with the standard deviation of 7 corresponded to intermediate-2 level of stop difficulty; and a non-smoothed image of ‘Y’ and ‘V’ letters (no 2-D Gaussian smoothing kernel filter applied) corresponded to high level of stop difficulty (hard ‘stop’ trials).

We implemented a dynamic staircase tracking procedure (Verbruggen & Logan, 2009) to individually adjust SSD based on a participant’s response to intermediate-1 level of stop difficulty. This was to ensure that the probability of successful inhibition (*P*(i)) was not affected by variability in the SSD but rather manipulated by the perceptual load of visual ‘go’ stimuli. After each successful ‘stop’ trial at intermediate-1 level of stop difficulty, the SSD was increased by 50ms making it harder for a participant to inhibit on subsequent ‘stop’ trials, and after each unsuccessful ‘stop’ trial at intermediate-1 level of stop difficulty, the SSD was decreased by 50ms, making it easier for a participant to stop on subsequent ‘stop’ trials. The SSD varied from 0-250ms.

On average, participants generated 354 correct ‘go’, 17 incorrect ‘go’, 76 correct ‘stop’, and 48 incorrect ‘stop’ trials (495 trials in total). Participants pressed ‘Q’ and ‘]’ button presses on the keyboard with their right and left index finger to make reaction time responses to ‘Y’ and ‘V’ letters. Stimulus-response maps were counterbalanced across participants.

### Data Analysis

On ‘go’ trials, the mean RT (ms) and the mean number of correct responses (%) were computed for each participant for each level of stop difficulty. On ‘stop’ trials, the mean stop-signal RT (SSRT) (ms) and the mean number of successful inhibitions *P*(i) (%) were estimated for each participant for each level of stop difficulty. We used the ‘integration method’ to calculate SSRT (ms) for each participant for each level of stop difficulty (Verbruggen & Logan, 2009). The correct ‘go’ RTs were ranked-ordered, and then the *n*th ‘go’ RT was selected, where *n* was estimated by multiplying the number of correct ‘go’ RTs in the distribution by the probability of responding (*P*(r)) to the visual ‘go’ stimuli at a given SSD (ms). We used one-way repeated measures ANOVA to examine whether variation in perceptual degradation of visual ‘go’ stimuli influence inhibitory processes in normal healthy individuals.

## Results

The mean RT (ms) and the mean number of correct responses (%) on ‘go’ trials, as well as the mean RT (ms) on ‘stop’ trials and the mean number of successful inhibitions *P*(i) (%), for each condition are shown in Table 1 and Figure 4.

**Fig. 4.**
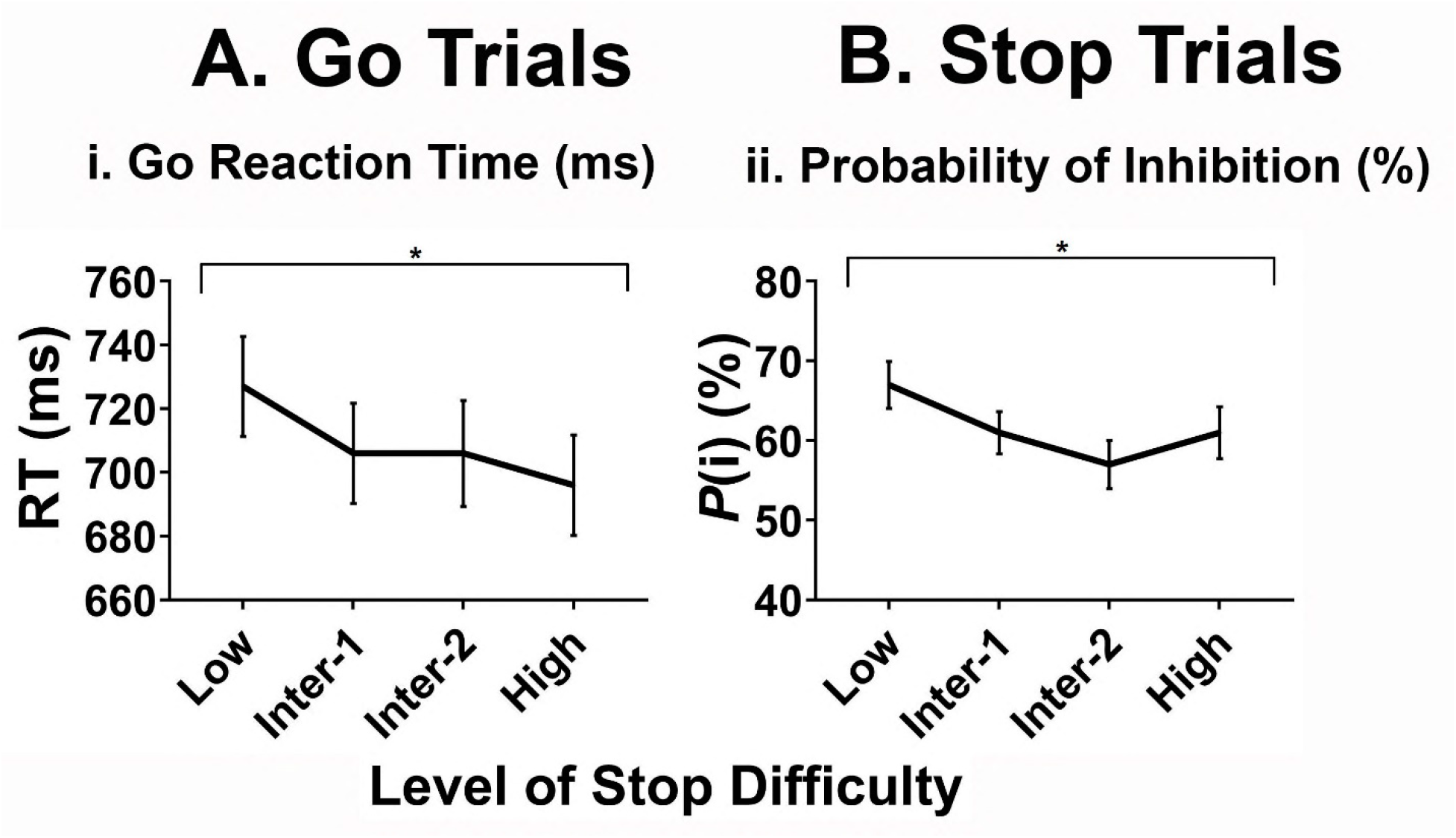
(A) Mean ‘go’ reaction times (ms) on ‘go’ trials for each level of stop difficulty. Standard errors are represented in the figure by the error bars for each stop difficulty level. Black asterisks (*) denote main effect of Level of Stop Difficulty on RTs for ‘go’ trials at *p* < .001. (B) Mean number of successful inhibitions *P*(i) (%) for each level of stop difficulty. Standard errors are represented by the error bars for each stop difficulty level. Black asterisks (*) denote main effect of Level of Stop Difficulty on *P*(i) for ‘stop’ trials at *p* < .001

**Table 1.** Behavioural Performance in normal healthy adults during modified stop-signal task.

| *Go Trials* | Low | Intermediate-1 | Intermediate-2 | High |
| --- | --- | --- | --- | --- |
| Means ± Standard deviations | | | | |
| Mean RT (ms) | 727.42 (79.90) | 705.73 (80.20) | 706.03 (84.92) | 695.96 (80.68) |
| Accuracy (%) | 94.81 (6.30) | 96.15 (4.81) | 95.85 (5.30) | 95.42 (5.29) |
| *Stop Trials* |  |  |  |  |
| $P(i)$ (%) | 66.70 (15.04) | 60.88 (13.53) | 56.81 (15.47) | 60.73 (16.63) |
| SSRT (ms) | 484.96 (43.52) | 485.27 (51.28) | 491.53 (48.30) | 468.31 (45.87) |
| Mean SSD (ms) | 160.94 (60.57) | 163.33 (52.09) | 175.29 (48.11) | 177.12 (48.51) |
| Mean Stopfailure RT(ms) | 623.50 (67.00) | 620.11 (90.50) | 626.50 (71.85) | 632.85 (70.80) |

### ‘Go’ Trials

Results from the one-way repeated measures ANOVA for ‘go’ RTs revealed a significant main effect of Level of Stop Difficulty, *F*(3, 75) = 23.84, *p* < .001, *ŋ^2^_p_* = .49 indicating that participants were faster with increased level of stop difficulty (see Figure 4(A)). This is expected because in the ‘higher’ stop difficulty conditions, the ‘go’ stimulus has little perceptual degradation and participants should take less time to identify it. Subsequent Bonferroni-adjusted pairwise comparisons (with an alpha level of .05/6 tests, a Bonferroni-correction at *p* = .0083) revealed that the ‘go’ RTs at low Level of Stop Difficulty (*M* = 727.42, *SD* = 79.90) were significantly slower than at intedmediate-1 (*M* = 705.73; *SD* = 80.20), intermediate-2 (*M* = 706.03; *SD* = 84.92) and high Levels of Stop Difficulty (*M* = 695.96; *SD* = 80.68), *p* < .001. Further, ‘go’ RTs at intermediate-2 (*M* = 706.03; *SD* = 84.92) of Stop Difficulty were slower than at high Level of Stop Difficulty (*M* = 695.96; *SD* = 80.68), *p* = .02. There was no significant main effect of Level of Stop Difficulty on the number of correct responses on ‘go’ trials, *F*(2.12; 52.91) = 2.04, *p* = .14, *ŋ^2^_p_* = .075.

### ‘Stop’ Trials

As expected, the one-way repeated measures ANOVA demonstrated that *P*(i) decreased as a function of Level of Stop Difficulty, *F*(3, 75) = 6.84, *p* <.001, *ŋ^2^_p_* = .22, which reflects that participants were less successful in inhibiting their response when the ‘stop’ trial was perceptually clear (hard ‘stop’ trials) (see Figure 4 (B)). Subsequent Bonferroni-adjusted pairwise comparisons (with an alpha level of .05/6 tests, a Bonferroni-correction at *p* = .0083) revealed that *P*(i) at low Level of Stop Difficulty (*M* = 66.70, *SD* = 15.04) was significantly higher than at intermediate-2 (*M* = 56.81; *SD* = 15.47) and high Levels of Stop Difficulty (*M* = 60.73, *SD* = 16.63), *p* < .001 and *p* = .031 respectively. There was no significant main effect of Level of Stop Difficulty on SSRTs, *F*(3, 75) = 2.61, *p* = .06, *ŋ^2^_p_* = .095 (this did not survive a Bonferroni-correction at *p* = .0083), suggesting that the time required to inhibit a response did not vary across levels of stop difficulty.

## Discussion and Conclusions

As expected, this study showed that with increased stop difficulty (little perceptual degradation), RTs on ‘go’ trials and the proportion of successful behavioural inhibitions on ‘stop’ trials (*P*(i)) decreased in normal healthy adults. There was no significant main effect of stop difficulty on the number of correct responses on ‘go’ trials and RTs on ‘stop’ trials. Importantly for our aim, the finding that participants were significantly less accurate in inhibiting motor responses with increased stop difficulty, indicates that stopping performance depends on the relative completion time of the ‘go’ and ‘stop’ processes, as predicted by the horse-race model. Overall, parametric manipulation of perceptual load of visual ‘go’ stimuli in the modified stop-signal task was partially successful.

Consistent with our findings, past studies have shown that perceptual task demands influence inhibitory control in normal healthy adults (Jahfari et al., 2013; Ma &Yu, 2016; Maguire et al., 2009). If we make visual ‘go’ stimuli perceptually challenging, individuals need additional time to process; this indirectly increases the relative completion time of the ‘go’ process, which results in successful stop-signal performance (slow ‘go’; ‘stop’ wins) (Schmidt et al., 2013; Verbruggen & Logan, 2009). However, on trials where visual ‘go’ stimuli are not subjected to perceptual degradation (fast ‘go’; hard ‘stop’), individuals process the stimuli more quickly, which is more likely to result in unsuccessful inhibition, as evidenced by decreased RTs on ‘go’ trials and lower *P*(i) on ‘stop’ trials. This finding highlights the important role of the completion time of the ‘go’ process in determining whether response inhibition is successful.

Contrary to our predictions, participants did not show increased SSRT with increased stop difficulty (fast ‘go’; hard ‘stop’). By contrast, there was a trend towards decreased SSRT (improved stop-signal performance) with increased stop difficulty, although this was not statistically significant. Stopping behaviour can be influenced by stop-signal intensity and modality (Pani et al., 2018), SSD length (D’Alberto et al., 2018; Hughes et al., 2013), and/or the complexity of visual ‘go’ stimuli (Groen et al., 2018; Yang, et al., 2014). Sensory and motor modalities rely on different sensorimotor neural pathways (Jahfari et al., 2015; Middlebrooks & Schall, 2014; Romei, Murray, Merabet, & Thut, 2007), and it is possible that the impact of stop difficulty on SSRT depends upon certain parameters of ‘go’ (face versus motion dots versus letters) and/or ‘stop’ stimuli (visual versus audio). For example, in a functional neuroimaging study, Jahfari et al. (2015) demonstrated that prefrontal-to-visual cortex connections may deliberately suppress visual input for the ‘go’ task, such that the inhibitory process can finish before the ‘go’ process, which results in shortened SSRT. As such, perceptual degradation of visual ‘go’ stimuli as a method for manipulating stop difficulty in the stop-signal paradigm may not be ideal, however, it is important to confirm whether certain stimulus and/or response modalities would differentially modulate *P*(i) and SSRT.

Our data further suggest that a race between direct ‘go’ and indirect ‘stop’ pathways is an incomplete account of basal ganglia involvement in motor control (Gu, Schmidt, & Berke, 2020; Mallet et al., 2016; Wei & Wang, 2016). Instead, the globus pallidus externus to striatum pathway can be critical for modulation of inhibitory processes in the cortico-basal ganglia circuitry (Hegeman, Hong, Hernandez, & Chan, 2016; Mallet et al., 2016; Wein & Wang, 2016). In a recent neurophysiological study by Mallet et al. (2016), ‘arkypallidal’ neurons in the globus pallidus externus were shown to selectively respond to ‘stop’ cue during stop-signal task thus signaling the cancellation of the action-in-preparation. These data suggest that the globus pallidus externus suppresses striatal ‘go’ process if required, and therefore is well-positioned to contribute to response inhibition (Gu et al., 2020; Mallet et al., 2016). Moreover, and according to an emerging view on the involvement of cortico-basal-ganglia loops in motor control, SSRT is modulated by the connectivity strength between different neural pathways (Wei & Wang, 2016). In particular, if the connection between the striatum and the substantia nigra increases (fast ‘go’ process), SSRTs increase, whereas if the globus pallidus externus-to-striatum connection gets stronger, SSRTs decrease (Wei & Wang, 2016). In line with that view, although participants in our study were less accurate in inhibiting motor responses with increased stop difficulty, it is possible that the fast ‘go’ process was not too fast to induce an increase in SSRT in normal healthy adults.

The findings of this study have improved our understanding of the interplay between response inhibition, perceptual degradation and task demands. We successfully manipulated the completion time of the ‘go’ process via perceptual degradation of visual ‘go’ stimuli, and found that increased stop difficulty influences inhibitory processes in two distinct ways. Specifically, while the fast ‘go’ process resulted in diminished number of successful behavioural stops in normal healthy adults, it did not increase SSRT, emphasising the possible role of other cortico-basal ganglia pathways, such as the globus pallidus externus-to-striatum, in addition to the striatum-to-substantia nigra, in modulating inhibitory processes.

In conclusion, we propose new insights into the relationship between the ‘go’ and ‘stop’ processes. Our findings imply that the horse-race model does not fully explain how different neural processes within the cortico-basal ganglia circuitry contribute to response inhibition. Further, we successfully used parametric perceptual degradation of visual ‘go’ stimuli to manipulate the relative completion time of the ‘go’ process, and provided evidence for the potential feasibility of this approach to modulate stop difficulty in the stop-signal paradigm – future neurophysiological evidence is warranted.

## Ethical statement

The study complies with the national ethical research guidelines in Australia. The study was approved by Monash University.

## Informed consent

Informed consent was obtained from each participant in accordance with the Helsinki Declaration.

## Acknowledgement

We acknowledge the financial support received from the Faculty of Science Postgraduate Publication Award.

Jamadar is supported by an Australian Research Council (ARC) Discovery Early Career Research Award (DE150100406) and the ARC Centre of Excellence for Integrative Brain Function (CE140100007).

We acknowledge the technical assistance of the National Imaging Facility, a National Collaborative Research Infrastructure Strategy (NCRIS) capability, at the Swinburne node.

We thank all the individuals for participating their time for this study.

